# Evaluation of methods for RNA-Seq analysis for uncovering key components of estrogen receptor-alpha signaling pathway in breast cancer

**DOI:** 10.1101/2024.03.16.585375

**Authors:** Wanru Guo

## Abstract

Breast cancer is the most common female cancer worldwide. Higher estrogen receptor (ER) expression is often associated with poor prognosis in ER positive breast cancer, however the exact mechanism is unknown. RNA-Seq data of three different experiments of ER knockdown (siE1, siE2, siE3) was used by researchers previously to identify TNFAIP1/BACURD2 as the mediator of ER induced increase in cell migration typical of breast cancer. We herein present a more comprehensive analysis of the data using DESEq2, along with comparison of results using a non-parametric approach, SAM-Seq, in order to cover the low sample size used in the study, and compared the results. We have found that, SAM-Seq uncovers more significant genes and is as robust as DESeq2 in discovering genes deemed significant by DESeq2. Excitingly, our approach was able to uncover three most significantly DE genes among the three independent experiments, namely, UHMK1, ACLY and CLIC4. The fact that they are involved in cancer regulation, metabolism and cell signaling, but so far has barely been studied, serves as additional exciting avenues of targeting ER pathway in breast cancer. Lastly, neither of our DESeq2 nor SAM-Seq analysis results showed consistent downregulation of TNPAIP1 upon ERR-alpha knockout across three independent experiments. Since our analysis approaches are both more conservative and robust in situations of model assumption violations, we came to the conclusion that further experiments are needed to ascertain the involvement of TNFAIP1 in the ER signaling pathway in breast cancer.

## 1. INTRODUCTION

### 1.1 Estrogen receptor signaling and role in breast cancer carcinogenesis

Breast cancer is the most common cancer amongst women in the US, it accounts around 30% (1 in 3) all new female cancer each year (American Cancer Society, 2024). In estrogen receptor (ER) positive breast cancers, the high expression is ERR-alpha correlates with tumor aggressiveness, in part through induced cell migration. In ER positive breast cancers, the high expression is ERR-alpha correlates with tumor aggressiveness through induced cell migration. Literature has shown has the Rho-GTPase, high RhoA expression reduces cell migration in breast cancer (Humphries, Wang, & Yang, 2020). However, upstream regulators are largely unknown and whether ERR targeting could affect this process is of interest to researchers. In the paper by Sailland *et al*., whether an ER mediated pathway affects RhoA stability therefore affecting cell migration is studied. Since the small GTPase RhoA inhibits actin thereby inhibiting cell migration, the hypothesis/aim by Sailland *et al*. (Sailland *et al*, 2014) was to find a mediator activated by ER that inhibits RhoA signaling, therefore promoting migration.

In the experiment by Sailland *et al*., RNA-seq of ER transcripts from three independent experiments of ERR-alpha knockdown using 3 siRNAs, siE1, siE2 and siE3 was conducted and compared to control, with two replicates in each condition, and total of 8 samples (n=8) with 23368 genes (p=23368). The paper found TNFAIP1 to be the mediator of cell motility by the ER receptor in estrogen-sensitive breast cancers. TNFAIP1 regulates RhoA stability by encoding BACURD2, a proteasome that degrades RhoA, therefore in turn triggering cell migration since RhoA acts to inhibit cell migration (Figure 1 as an illustration).

**Fig 1.**
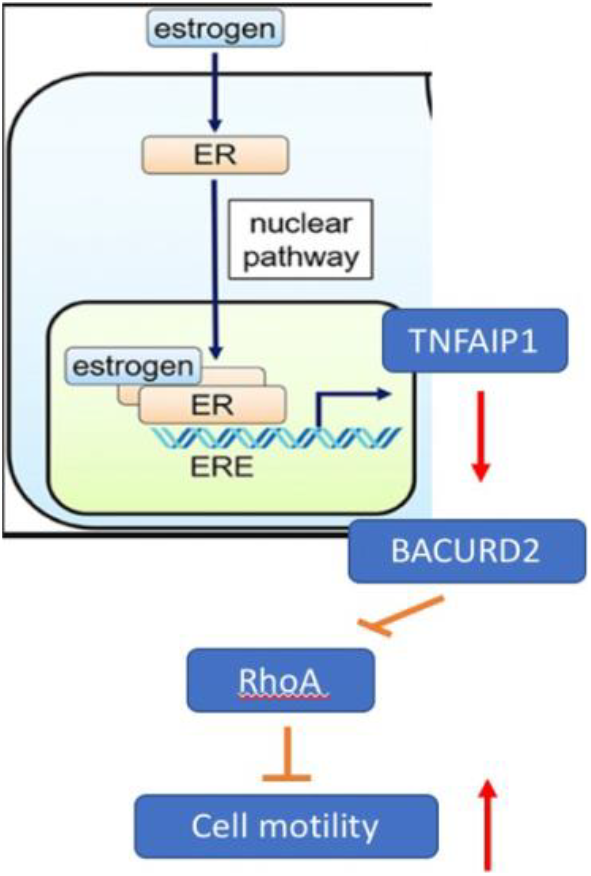
The ER pathway suggested by Sailland *et al*. on promoting cell motility and tumor aggressiveness in breast cancer.

### 1.2 RNA-Seq analysis using DESeq2

In the analysis by Sailland *et al*., genes are deemed to be upregulated/downregulated by looking at their fold changes (>1.5 and <0.75 respectively), and declared significance of TNFAIP1 using three replicates (since n=3 is enough to declare a finding in biology). However, we are interested in analysis of the data from a statistical perspective. In common RNA-Seq analysis methods such as DESeq2, the significance of genes are determined using Z-score of the Wald statistic, which is calculated taking into consideration both the logFC and its standard error, namely, 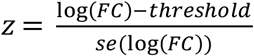, which in turn calculates the false discovery rate (adjusted p-value). Merely relying on log fold changes in determining which genes are up- and down-regulated, without consideration the standard error, could lead to incorrect conclusions regarding significance. Therefore, in our analysis, we will re-examine the significance of TNFAIP1.

### 1.3 Non-parametric analysis using SAM-Seq

The parametric methods, DESeq2, models the count of each gene using a negative binomial (NB) distribution, where the variances are proportional to the means squared. However, this assumption does not always hold, such as in the case when the sample size is small (in our study, n=2 for each condition), or if there are outliers present (not applicable in our study since we only have n=2 for each condition). In this case with low sample size, we reasoned that a non-parametric method, such as SAM-Seq would be useful, which has been shown to perform competitively with DESeq2 under situations where the assumption holds, and outperforms DESeq2 when the assumption does not (Li & Tibshirani, 2013). The SAM-Seq procedure is as follows. Suppose we have a N x P matrix for expression count data, where N= No. of samples and p = No. of features (genes). For Feature *j*, suppose that we have counts *N*_1*j*_, …, *N*_*nj*_ from either Class 1 or Class 2. n=4 there are 4 total samples, *k* = 1, 2, where n1 = n2 = 2, and *n*1 + *n*2 = 4. Let *Ck* = {*i* : Sample *i* is from Class *k*}, *k* = 1, 2. Let *R*_*ij*_(*N*) be the rank of *N*_*ij*_ in *N*_1*j*_, …, *N*_*nj*_. Then, the two-sample Wilcoxon score statistic is:

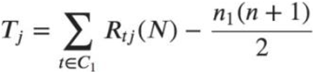

It then compares the observed score distribution with the null distribution, and then computes FDR by permutation plug-in method typical in the non-parametric setting using 100 permutations. Because of its utilization of score statistic based on rank, it yields the same score, and hence FDR, regardless of outliers or low (zero) counts; whereas a parametric method, taking the average of the expression in samples in the same condition, are more affected by outliers/ zero or low counts.

Given our small sample size, it would be interesting to evaluate its performance against the parametric method, DESeq2. We could compare the FDR, number of significant genes called, and especially the significance of the gene TNFAIP1, as well as analyze the characteristics of the differences. We hypothesize that given our small sample size (2 replicates each condition), the parametric assumption may not hold, so we would expect SAM-Seq to be more robust than DESeq2, and uncover more significant genes.

## 2. MATERIALS AND METHODS

In order for our method to call approx. the same number of DE genes as in Sailland *et al*., we applied a lfc threshold of 0.32 adjustment in our analysis. This would make sure we use the correct method of analysis (FDR and not the log-fold change as a measure of significance), and at the same time not having too many significant genes. DESeq2 was performed for 3 independent experiments using siE1, siE2 and siE3 knockdown of ERRalpha using the Bioconductor R package, and logFC for each gene is computed in the software with the wald statistic generated using 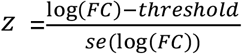 to obtain the p-values, after which the BH method for multiple testing was used to obtain the FDR (Padj). Genes are called significant DE genes if FDR (Padj) < 0.05.

We also analyzed the RNA-Seq data using SAM-Seq from the R package samr. We firstly compared SAM-seq with DESeq2 in the analysis of siE1 knockdown experiment without applying the lfc threshold for ease of comparison of the performance of the two methods. The procedure of SAM-Seq was described in the Introduction section. In short, we use a Wilconxin T statistic which is based on the sum of the ranking of the samples for each gene, and then applied permutation to compare our distribution of the statistic with that of the expected null distribution, where FDR is computed using the permutation plug-in approach. We compared the two methods in terms of the overall FDR, the number of significant genes, and the characteristics of genes deemed significant (FDR < 0.05) by SAM-Seq but not called by DESeq2, and correlated the significant genes called by SAM-Seq with that called by DESeq2.

Lastly, we applied SAM-Seq2 to the analysis of entire dataset (siE1,siE2,siE3) and compared the significance result of TNFAIP1 with that of DESeq2.

## 3. RESULTS

We conducted an analysis using DESeq2 employing wald-test based FDR as opposed to the log-fold change approach presented previously. We found that we obtained much more genes, making comparison difficult. Therefore, a lfc threshold of 0.32 was applied in our analysis, and we yielded 1260, 798 and 1736 significant DE genes independently modulated by siE1, siE2 and siE3 experiments. We found 202 genes in common that were regulated all three experiments (Fig. 2). The volcano plots showing the significant genes in terms of FDR are labeled in green (Fig. 3). This method ensures that we obtain comparable number of significant genes in our analysis, while at the same time employing FDR to call significant genes, not fold changes.

**Fig 2.**
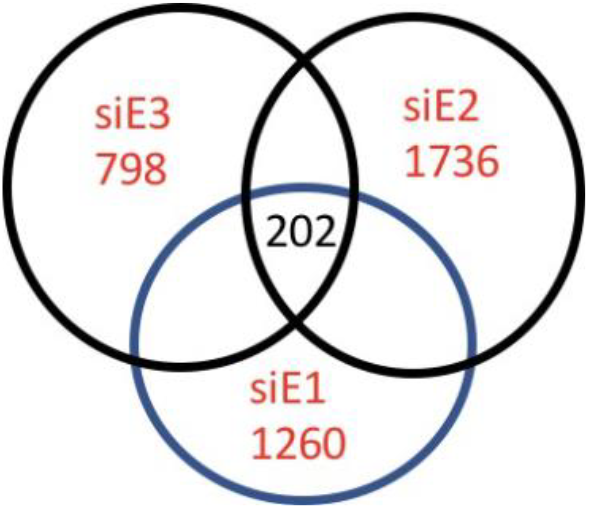
DEseq2 analysis of breast cancer RNA-Seq data on three ER siRNA knockdown experiments

**Fig 3.**
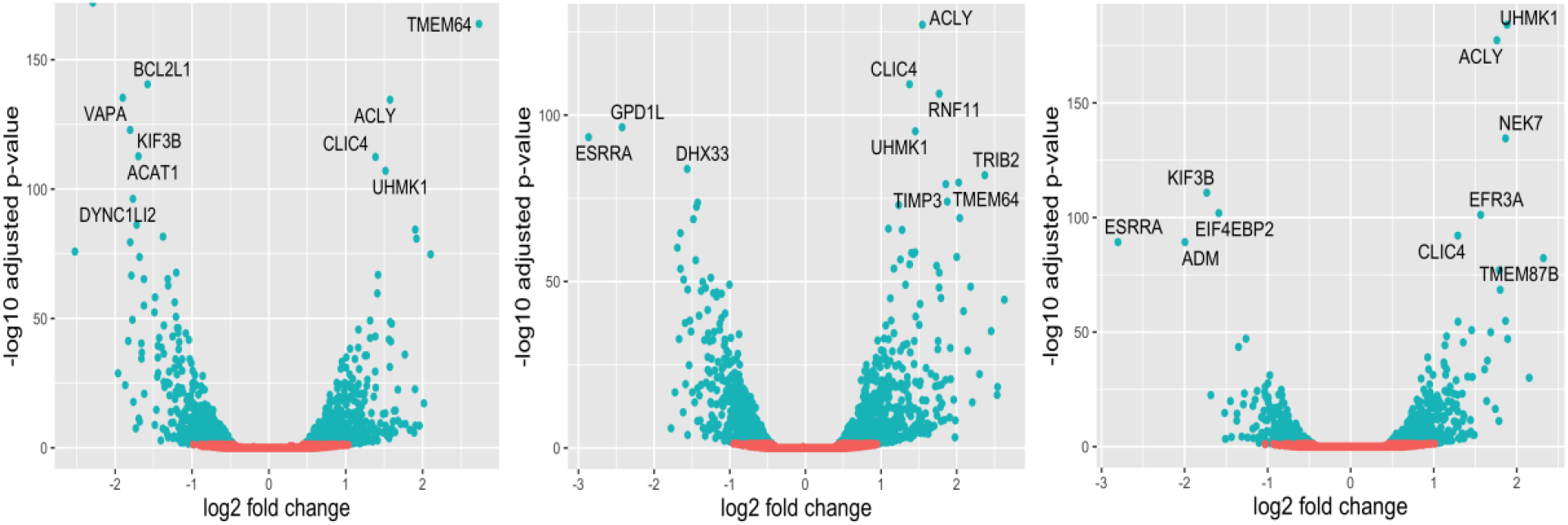
Genes regulated by three ER siRNA knockdown experiments analyzed using DESeq2, siE1 (left), siE2 (middle) and siE3 (right) (DE genes are labelled in green with FDR threshold <0.05)

### 3.1 DESeq2 analysis also uncovers important DE genes (UHMK1, ACLY, CLIC4) involved in cancer pathway

We thereafter compared the top 10 significant genes uncovered by our analysis across the experiments siE1, siE2 and siE3. We found that three genes, UHMK1, ACLY, and CLIC4 are shared by the 3 experiments as top 10 genes (Table 1). After performing literature search, we found that they are all involved in the cancer pathway, namely, UHMK1 upregulated in gastric cancer through reprogramming nucleotide metabolism (Feng *et al*., 2020); CLIC4 is associated with NFkB signaling pathway (Suh *et al*., 2012); and ATP citrate Lyase (ACLY), activates of Akt signaling and promotes proliferation (Khwairakpam *et al*., 2015). This presents an exciting avenue as targets of breast cancer mediated by the ER pathway, that is discovered by our analysis.

**TABLE 1.**
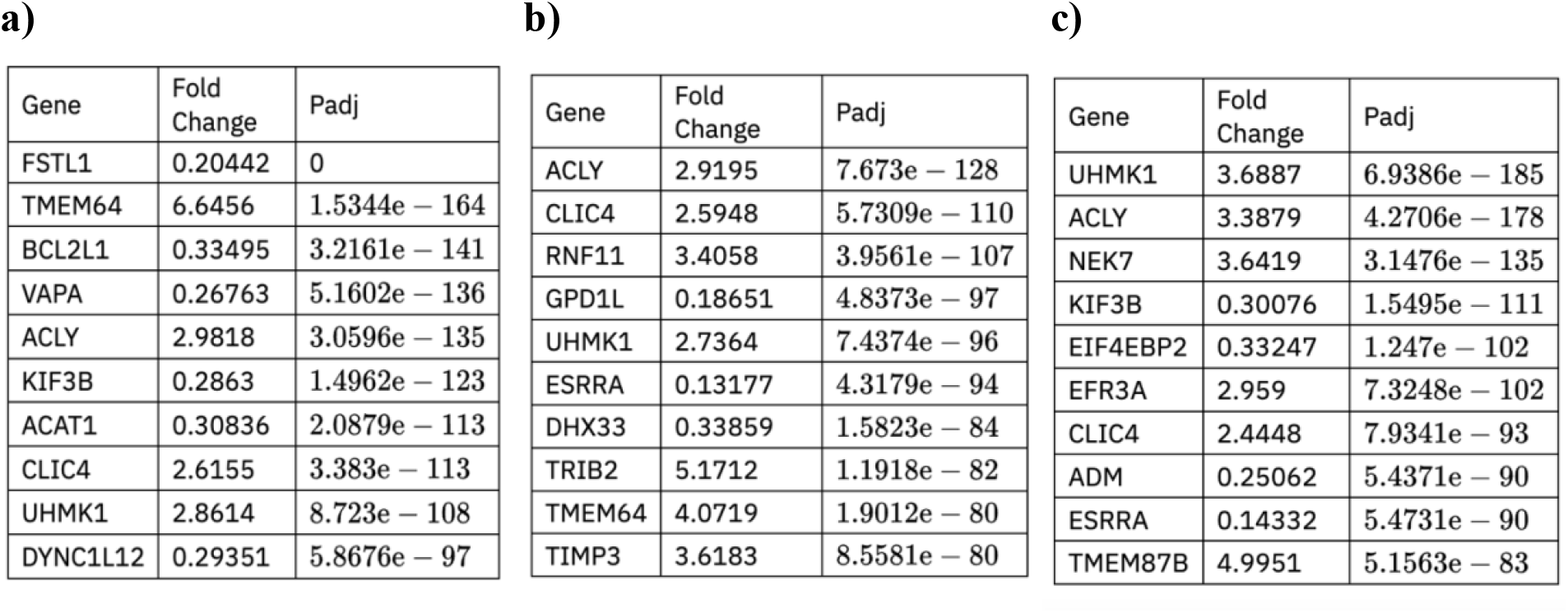
Top 10 genes uncovered by Deseq2 reanalysis of a) siE1, b) siE2, and c) siE3 KD of ERR experiments.

### 3.2 TNFAIP1 is not consistently downregulated by ERR in three independent experiments in our analysis

Given the critical role of TNFAIP1 in ER-mediated breast cancer regulation, we would like to examine the significance of TNFAIP1 in our analysis. In table 2, fold changes and FDR of the gene TNFAIP1 in three independent experiments are shown, along with the result yielded by Sailland *et al*. We that see that the fold changes are the same, which is expected since the KD of ERR-alpha downregulates TNFAIP1, consistent with the theory in the paper that TNFAIP1 transcription is activated by ERR-alpha signaling in breast cancer (Sailland *et al*, 2014).

**TABLE 2.** Comparison of TNFAIP1 expression in 3 ERR knockdown experiments in a) Sailland *et al*. and b) our DESeq2 analysis.

| Experiment | Fold Change | Padj |
| --- | --- | --- |
| siE1 | 0.57 | $1.80e - 23$ |
| siE2 | 0.49 | $1.25e - 35$ |
| siE3 | 0.73 | $1.93e - 05$ |

b)
| Experiment | Fold Change | Padj |
| --- | --- | --- |
| siE1 | 0.57214 | 0.0000020995 |
| siE2 | 0.50379 | $9.3206e - 12$ |
| siE3 | 0.73727 | 1 |

However, siE3 did not yield significant results for TNFAIP1 in our analysis (FDR = 1); it was however deemed significant by Sailland *et al* (padj = 1.93e-5), This finding is interesting since it shows that the gene TNFAIP1 may not be consistently differentially downregulated in ERR knockdown across 3 independent experiments, meaning that more experiments are needed to confer biological plausibility and to ascertain TNFAIP1 as a target for ER signaling.

### 3.3. SAM-Seq identified more significant genes than DESeq2, even though FDR is inflated in the significant genes identified by SAM-Seq

Next, we sought to compare the performance of our DESeq2 method with the non-parametric method, SAM-Seq. This is motivated by the fact that our sample size is small (n=2 per condition, and n=8 total), thereby may not follow parametric assumptions, and employing the non-parametric method may uncover significant genes not discovered by DESeq2. We utilized a single experiment, siE1 to compare the methods, where DESeq2 without lfc threshold adjustment was used for comparability. We compared the FDR for the 10,000 most significant genes.

Firstly, we found that SAM-Seq called 7470 genes as significant, while DESeq2 called 5129. Secondly, Amongst the significant genes, the FDR for SAM-Seq is higher than DESeq2, even though the overall FDR is lower (Figure 4). This is expected, since SAM-Seq computes FDR according to permutation plug-in method, which compares the observed score distribution with the null distribution, thereby inflating the FDR. We also see from Fig. 3 that many FDR values in SAM-Seq are tied, mainly because FDRs are determined using Wilcoxin scores based on ranking of the count across samples, which does not distinguish between the actual count values. This results in tied scores and thus tied FDR values.

**Fig 4.**
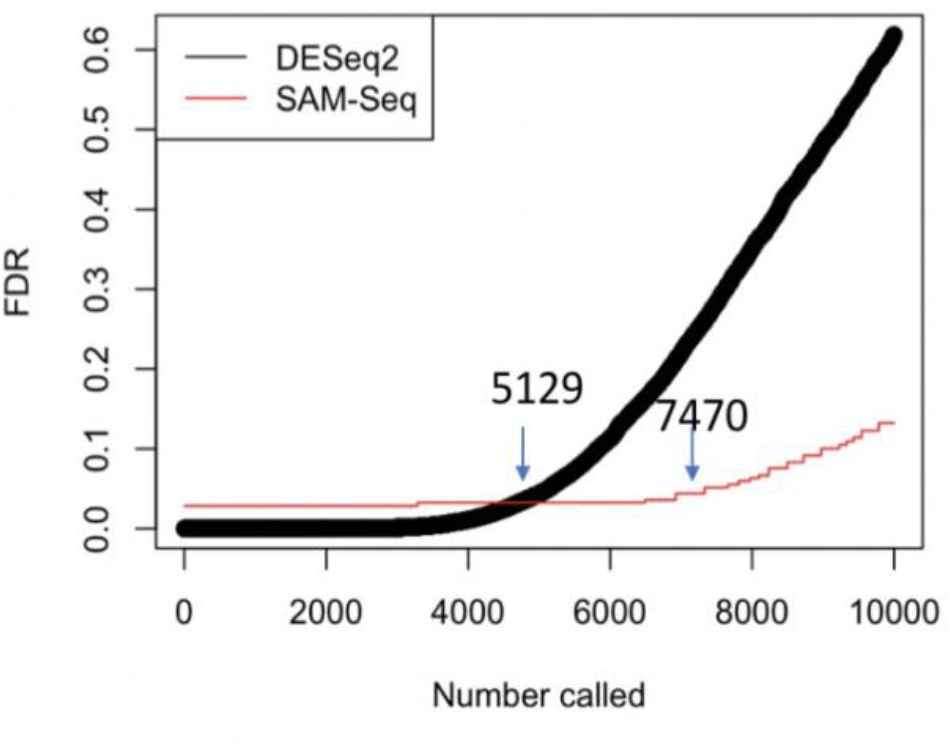
FDR values for the top 10,000 most significant genes for DESeq2 and SAM-Seq in the siE1 experiment. Using 0.05 as the significance threshold, DESEq2 uncovered 5129 genes, while SAM-Seq discovered 7470.

### 3.4 SAM-seq uncovers potentially differentially expressed genes deemed non-significant by DESeq2

We next wondered what characteristics makes the gene called significant by SAM-Seq, but not significant by DESeq2. We selected 4 genes (Table 3) that fall into this definition from a single experiment siE1. We found that all genes share a common characteristic – their read counts are very low (VWCE and TMEM240) or even zeros (TGM5 and TBC1D10C) (Table 3). Since the sample size is small (n=2 each condition), the parametric method NB model would preferentially call genes with higher read counts significant, even if the absolute fold changes are actually similar. According to table 3, we see that although the absolute fold changes for these 4 genes are quite big (around 4-5 fold for VWCE and TMEM240), the fold changes determined by the NB model are 1.25 and 1.24 respectively. This is because these low counts (especially zero counts) do not fit the regular NB model well, where a zero-inflated model should be used instead.

**TABLE 3.** Expression patterns of genes that are significant by SAM-Seq but insignificant by DESeq2.

| Gene | SAM-Seq Score | FDR | DESeq2 Fold Change | p-value | Control 1 counts | Control 2 counts | siE1 – 1 counts | siE1 – 2 counts |
| --- | --- | --- | --- | --- | --- | --- | --- | --- |
| VWCE | 2 | 0.03026 | 1.2525 | 0.11722 | 2 | 3 | 10 | 9 |
| TMEM240 | 2 | 0.03026 | 1.2360 | 0.12102 | 2 | 2 | 7 | 10 |
| TGM5 | 2 | 0.03026 | 1.1490 | 0.05934 | 0 | 0 | 4 | 4 |
| TBC1D10C | 2 | 0.03026 | 1.0927 | 0.16956 | 0 | 0 | 3 | 2 |

However, a non-parametric Wilcoxin rank score does not depend on the actual absolute counts of the gene, instead, only their ranking across samples within each gene matter. Therefore, there could be a situation where for genes A and B, the absolute fold changes (and LogFC) are the same, however, gene A had lower read counts than gene B therefore not called significant by DESeq2. However, SAM-Seq is still able to call gene A significant since it does not rely on absolute counts. This is a very useful way of uncovering genes that are potentially truly DE genes, but are missed because of their low counts. In this case in the dataset, the genes VWCE, TMEM240, TGM5 and TBC1D10C could all be potentially significant DE genes but are missed. On the other hand, we should also be wary of this method since it might also generate false positives, since we are using real datasets and do not know which genes are truly DE genes.

### 3.5 SAM-Seq is robust in identifying significant genes called significant in DESeq2

Now that we found that SAM-Seq is robust in identifying potentially significant genes that are missed by DESeq2. However, for significant genes called by DESeq2, could SAM-Seq also identify them? We answer this question using 2x2 Chi-squared test to examine the association between the genes deemed significant by DESeq2 and those deemed significant by SAM-Seq (Table 4). We found significant association between significant genes called by the two methods, with Chi-squared statistic of 13845, and p-value < 2e-16. This means that for differentially expressed genes by DESeq2, SAM-Seq would also deem them significant. This is evident in table 4, as 5129 genes significant by DESeq2, of which 5112 also significant in SAM-Seq; we also found 18239 genes non-significant by DESeq2, of which, 15881 also deemed non-significant by SAM-Seq, therefore, we can see that SAM-Seq is able call almost all of the significant gene by DESeq2 significant. This means that SAM-Seq is not only able to uncover potentially significant genes missed by DESeq2, it is also able to call the same genes deemed significant by DESeq2 significant.

**TABLE 4.** 2X2 Chi-Squared table of gene significance by DESeq2 and SAM-Seq.

| DESeq2 | SAM-Seq |  |
| --- | --- | --- |
|  | Significant | Non-significant |
| Significant | 5112 | 17 |
| Non-significant | 2358 | 15881 |

### 3.6 TNFAIP1 is also not consistently significant across experiments through analysis using SAM-Seq

We finally asked ourselves, would the lack of significance of TNFAIP1 in siE3 experiment be the result of poor fit NB model to the data with low sample size, therefore making the log fold change (0.73) appear milder than the actual effect, *or does this experiment really yield no significance?* Answering this question is important because it determines whether we should repeat the study at least several more times to ascertain the involvement of TNFAIP1 in breast cancer. To do this, we employ our non-parametric method SAM-Seq which is shown (Sections 1.2 and 1.3) to be robust against situations that deviate from NB distribution. As table 5 has shown, interestingly, siE3 experiment re-analysis by SAM-Seq also yielded non-significant FDR (0.06), which is consistent with our DESeq2 analysis done previously. The other experiments, siE1 and siE2 all generated significant results, also consistent with our previous findings using DESeq2. Therefore, we can conclude that we indeed could not find a biologically significant change and not due to the structure of our data/low sample size, which means that the consistency of the findings that TNFAIP1 is downregulated upon ERR knockdown is questionable, and more future experiments indeed need to be conducted to impose biological reproducibility.

**TABLE 5.** TNFAIP1 differential expression in 3 independent experiments (ERR knockdown) analyzed by SAM-Seq.

| Experiment | Fold Change | FDR |
| --- | --- | --- |
| siE1 | 0.569 | 0.03060 |
| siE2 | 0.504 | 0.03092 |
| siE3 | 0.74 | 0.06287 |

## 4. DISCUSSIONS

Breast cancer as the most common cancer amongst females in the US, where a good proportion is estrogen-receptor positive, the exact mechanism through which ER signals to promote cell migration is largely unknown and targeting this pathway would present a promising therapeutic strategy. Since ER is a nuclear receptor (Paterni *et al*, 2014) and acts as transcription factor itself, RNA-Seq analysis of ER transcripts are crucial in revealing important information, far surpassing traditional molecular biology approaches such as qPCR or western blots in that it is high-throughput. The findings by Sailland *et al*. that ER signaling to promote migration through TNFAIP1, which encodes BACURD2 and degrades RhoA is exciting. However, employing our methods on DESeq2 and SAM-Seq as detailed in this paper reveals that the downregulation of TNFAIP1 upon ERR knockout, which is found in Sailland *et al*. yielded inconsistent results across 3 independent experiments, may require further research, as one of the experiments is deemed insignificant by both of our analyses (DESeq2 and SAM-Seq). This inconsistency however, is not due to the lower sample size or structure of the data, since our non-parametric method, SAM-Seq, which is found by our study to be more robust than the parametric method, DESeq2, also failed to obtain a significant result for the differential regulation of TNFAIP1 by ERR KD in that particular experiment. Together, this warrants further experiments to be done to reach a biologically meaningful conclusion. Since n=3 is the minimum number of independent experiments needed to reach a plausible conclusion in biology, more ERR KD experiments *with increased sample size* are needed to ascertain the involvement of TNFAIP1 in ER/RhoA pathway leading to enhanced cell motility in breast cancer.

Another significant finding in our study is that SAM-Seq could uncover potentially DE genes deemed insignificant by DESeq2 because of their low counts or presence of zero counts, while performing as powerful as DESeq2 in discovering genes deemed significant by DESeq2. This exciting finding from our data means that one no longer needs to worry about the failure of RNA-Seq experiments to meet parametric assumptions, since at times low sample size, outliers, zero or low counts are inevitable especially when there are time and resources constraints.

However, one important consideration here is we do not actually if the additional genes uncovered by SAM-Seq are truly DE genes or not, it may be due to chance that a difference is observed simply because of the low count of the data. Hence, we could not truly evaluate the performance of these two methods since we used real datasets of ER transcriptomics. Future studies using simulated datasets would be ideal to compare the sensitivity and specificity of these methods (e.g. AUC) to know how well exactly they work under situations where the parametric assumptions are met and not met, therefore reaching meaningful conclusion that could in future be applied to datasets displaying similar characteristics. Only then would we know under what situations would SAM-Seq outperform DESeq2.

There are several limitations of the study using SAM-Seq, though, besides the fact that the true DE genes are not known, firstly, since FDRs are calculated from permutation plug-in method, it could be less accurate than parametric methods, and therefore the real significance as determined by the FDR threshold of 0.05 is questionable, as we saw that most FDRs are more inflated compared to DESeq2 especially for significant genes. Secondly, probably the most important point is that, because the FDR is calculated using the permutation method, it is not reproducible across studies, therefore not replicable. This could be a huge problem though when it comes to biologically meaningful conclusions, such as in this case with the role of TNFAIP1 in breast cancer ER signaling. Thirdly, since the non-parametric methods works based upon rank statistic (Wilcoxin statistic), this results in the T scores/FDRs of a large portion of genes to be indistinguishable from another, in this experiment, more than 3000 genes have the same FDR of 0.03026, even though their count data distribution are very different. DESeq2, on the other hand, distinguishes each gene based on subtle differences in their expression count data and generates FDR that easily enables ranking of these genes in terms of e.g. top 10 significant genes, whereas finding the top 10 significant genes is almost impossible using SAM-Seq. Therefore, due to these drawbacks, we should only really reserve the use of non-parametric methods such as SAM-Seq to situations where we have observed the data and identified that parametric assumptions are not met, otherwise DESeq2 would still be preferred.

It has also been brought to our attention that, although our data is suitable for evaluating the performance of non-parametric model (vs. parametric model) under limited sample size and inflated zeros, it is unsuitable for examining the effect of outliers on model performance – an important scenario shown in literature (Li & Tibshirani, 2013) where SAM-Seq would outperform DESeq2. In order to examine the effect of outliers, we need to repeat the experiment utilizing samples where there are for example, 10 or more replicates for each condition (e.g. siRNA E1 and control), and compare situations where one count within each condition is much different from others (outlier scenario), with those with consistent expression across samples in the same condition (without outliers). We hypothesize that SAM-Seq is shown to perform better than DESeq2 when outliers are present, giving more accurate/reliable results than DESeq2, which is because the rank based statistic are less affected by outliers, where for DESeq2, one outlier would alter the significance of the gene since DESeq2 relies on the average in each condition, resulting in either false positives or failure to discover significance for true DE genes. Taking this point into consideration, in the future, when in the future, reconducting the siRNA experiments for ER knockout to replicate the findings of TNFAIP1 downregulation, apart from using increased sample size conducting more independent experiments, we would also be wary of outliers in the data. If such outliers are present, the non-parametric method SAM-Seq should be used to give a more accurate estimate of the true significance of TNFAIP1. If our data meet parametric assumptions, we would still use DESeq2 in our analysis, due to the several aforementioned drawbacks of SAM-Seq, including inflation of FDR and inability to rank the genes in terms of significance and discover the most significant genes. If we are able reach more than three consistently significant results using our improved experimental design and statistical analysis, we could truly declare the ER/TNFAIP1/BACURD pathway as a promising therapeutic for ER positive breast cancer, which is an exciting prospect.

Last, but not least, the fact in our 3 independent experiments we found that amongst the top 10 significant gene discovered using this conservative approach, 3 of them are commonly regulated, namely, UHMK1, ACLY and CLIC4 really gives us confidence that these genes are indeed modulated by ER signaling in breast cancer as a very conservative and stringent approach is used in our study. Upon a literature search, these genes are indeed involved in cancer pathways. UHMK1 upregulated and promotes gastric cancer through reprogramming nucleotide metabolism (Feng et al., 2020); CLIC4 is found to be associated with Arf6 mediated NFkB signalling pathway in AML (Suh et al,, 2012); ATP citrate Lyase (ACLY), those activation catalyzes the conversion of citrate to oxaloacetic acid (OAA) and acetyl-CoA, and in cancer leads to increased metabolic activity via activation of Akt signaling and promotes proliferation (Khwairakpam et al., 2015). However, there’s very little evidence that these genes are involved in ER signaling in breast cancer according to literature. Therefore, our study would also serve to uncover additional mediators of ER signaling pathway, that, albeit unrelated to Rho-GTPase binding and cell migration, could still present as exciting new targets in breast cancer treatment that involves regulating the cell cycle/metabolism.

## 5. CONCLUSIONS

A more accurate and conservative approach using parametric method using DESeq2, and non-parametric method, SAM-Seq, was used to re-analyze ER knockdown expression data by 3 independent experiments (siE1, SiE2, siE3). SAM-Seq, a non-parametric method, could uncover potentially DE genes deemed insignificant by DESeq2 because of their low counts or presence of zero counts, while at the same time, for genes that are discovered as DE genes by DESeq2, SAM-Seq could also call/discover them. Our analysis of TNFAIP1 expression showed that both the parametric method DESeq2, and non-parametric method, SAM-Seq which is shown to be more robust by our study and others (Li & Tibshirani, 2013), both fail to give significant results for siE3 experiment. Since n=3 is the minimum number of independent experiments needed to reach a plausible conclusion in biology, more ERR KD experiments with increased sample size are needed to ascertain the involvement of TNFAIP1 in ER/RhoA pathway leading to enhanced cell migration in breast cancer. Even though SAM-Seq should be used when parametric assumptions are not met, DESeq2 is still the preferred method of analysis when the data distribution follows the parametric assumption. This is because of several drawbacks discovered in our study, including lack of replicability and inflated FDRs, and many tied FDR values making discovering the top most significant DE genes impossible. Lastly, our analysis in 3 independent KD experiments uncovered 3 common DE genes amongst the top 10 most significant genes, UHMK1, CLIC4 and ACLY. The fact that they are involved in cancer regulation upon literature search, but so far has barely been studied, serves as additional exciting avenues of targeting ER pathway in breast cancer.

## ETHICS APPROVAL AND CONSENT TO PARTICIPATE

Not applicable.

## CONSENT FOR PUBLICATION

Not applicable.

## AVAILABILITY OF DATA

The RNA-seq data are available in the Gene Expression Omnibus database (www.ncbi.nlm.nih.gov/geo/) with Accession No. GSE49110.

## ACKNOWLEDGEMENTS

I am profoundly grateful for the support and resources that have been available to me throughout the development of this paper. Special thanks to the Biostatistics Department’s staff at my alma mater, whose open-door policy allowed me to access office space. I would also like to acknowledge the constructive feedback provided by my peer reviewers, who offered critical insights that greatly enhanced the quality of this manuscript.

My gratitude extends to my family for their understanding and patience throughout the countless hours spent on this project.

## REFERENCES

[1] American Cancer Society. Breast cancer statistics: How common is breast cancer? American Cancer Society. 2024. Accessed Mar 10, 2024. https://www.cancer.org/cancer/types/breast-cancer/about/how-common-is-breast-cancer.html.

[2] Feng, X., Ma, D., Zhao, J., Song, Y., Zhu, Y., Zhou, Q., Ma, F., Liu, X., Zhong, M., Liu, Y., Xiong, Y., Qiu, X., Zhang, Z., Zhang, H., Zhao, Y., Zhang, K., Hong, X., & Zhang, Z. UHMK1 promotes gastric cancer progression through reprogramming nucleotide metabolism. EMBO J. 2020; 39(5): e102541. 10.15252/embj.2019102541

[3] Humphries, B., Wang, Z., & Yang, C. (2020). Rho GTPases: Big Players in Breast Cancer Initiation, Metastasis and Therapeutic Responses. Cells, 9(10), 2167. 10.3390/cells9102167

[4] Khwairakpam, A.D., Shyamananda, M.S., Sailo, B.L., Rathnakaram, S.R., Padmavathi, G., Kotoky, J., & Kunnumakkara, A.B. ATP citrate lyase (ACLY): a promising target for cancer prevention and treatment. Curr Drug Targets. 2015; 16(2): 156–163. 10.2174/1389450115666141224125117

[5] Li, J. & Tibshirani, R. Finding consistent patterns: a nonparametric approach for identifying differential expression in RNA-Seq data. Stat Methods Med Res. 2013; 22(5): 519–536. 10.1177/0962280211428386

[6] Paterni, I., Granchi, C., Katzenellenbogen, J. A., & Minutolo, F. (2014). Estrogen receptors alpha (ERα) and beta (ERβ): subtype-selective ligands and clinical potential. Steroids, 90, 13–29. 10.1016/j.steroids.2014.06.012

[7] Sailland, J., Tribollet, V., Forcet, C., Billon, C., Barenton, B., Carnesecchi, J., Bachmann, A., Gauthier, K.C., Yu, S., Giguère, V., Chan, F.L., & Vanacker, J.M. Estrogen-related receptor α decreases RHOA stability to induce orientated cell migration. Proc Natl Acad Sci USA. 2014; 111(42): 15108–15113. 10.1073/pnas.1402094111

[8] Suh, K.S., Malik, M., Shukla, A., Ryscavage, A., Wright, L., Jividen, K., Crutchley, J.M., Dumont, R.A., Fernandez-Salas, E., Webster, J.D., Simpson, R.M., & Yuspa, S.H. CLIC4 is a tumor suppressor for cutaneous squamous cell cancer. Carcinogenesis. 2012; 33(5): 986–995. 10.1093/carcin/bgs11

